# Common species link global ecosystems to climate change

**DOI:** 10.1101/043729

**Authors:** Bjarte Hannisdal, Kristian Agasøster Haaga, Trond Reitan, David Diego, Lee Hsiang Liow

## Abstract

Common species shape the world around us, and changes in their commonness signify large-scale shifts in ecosystem structure and function^1-4^. Dominant taxa drive productivity and biogeochemical cycling, in direct interaction with abiotic components of the Earth system^3,4^. However, our understanding of the dynamic response of ecosystems to global environmental changes in the past is limited by our ability to robustly estimate fossil taxonomic richness^5,6^, and by our neglect of the importance of common species. To rectify this, we use observations of the most common and widespread species to track global changes in their distribution in the deep geological past. Our simple approach is robust to factors that bias richness estimators, including widely used sampling-standardization methods^5^, which we show are highly sensitive to variability in the species-abundance distribution. Causal analyses of common species frequency in the deep-sea sedimentary record detect a lagged response in the ecological prominence of planktonic foraminifera to oceanographic changes captured by deep-ocean temperature records over the last 65 million years, encompassing one of Earth's major climate transitions. Our results demonstrate that common species can act as tracers of a past global ecosystem and its response to physical changes in Earth's dynamic history.

---

True species richness can be elusive even in well-studied ecosystems, because most species are very rare, and relatively few species account for most of the total abundance^1,2^. For example, only ~1.4 *%* of the estimated tree species account for half of the biomass and control the cycling of water, carbon, and nutrients in the Amazon forest^3^. Similarly, a recent survey of eukaryotic diversity in the oceans found that ~0.24 % of the taxa accounted for half of the total number of rDNA reads^4^.

Abundance and occupancy are typically positively correlated, with the more abundant species being the more widespread^4,7^. In the fossil record, species and higher taxa generally have a humped temporal distribution of occurrences, being rare in the early and late stages of their known stratigraphic range^8-10^.

Here we accommodate these ecological features by focussing only on species that are common and widespread at any given time, using the Summed Common species Occurrence Rate (SCOR), a very simple occurrence-based quantity that is sensitive to changes in total abundance (Methods)^11^. We apply SCOR to deep-sea sedimentary records of calcifying plankton (coccolithophores and foraminifera) over the last 65 years to demonstrate how relative changes in the distribution of common and widespread species were linked to climate change on geological time scales.

First, we evaluate the sensitivity of SCOR and commonly used richness estimators to potential biases using Poseidon, a simulation model of planktonic microfossil occurrences (Fig 1; Methods and Supplementary Code). We target methods currently popular in palaeobiology, and highlight the effects of two main factors: variability in the spatial sampling completeness (Fig. 1b), and variability in the shape of the species rank-abundance distribution (RAD; Fig. 1c).

**Figure 1.**
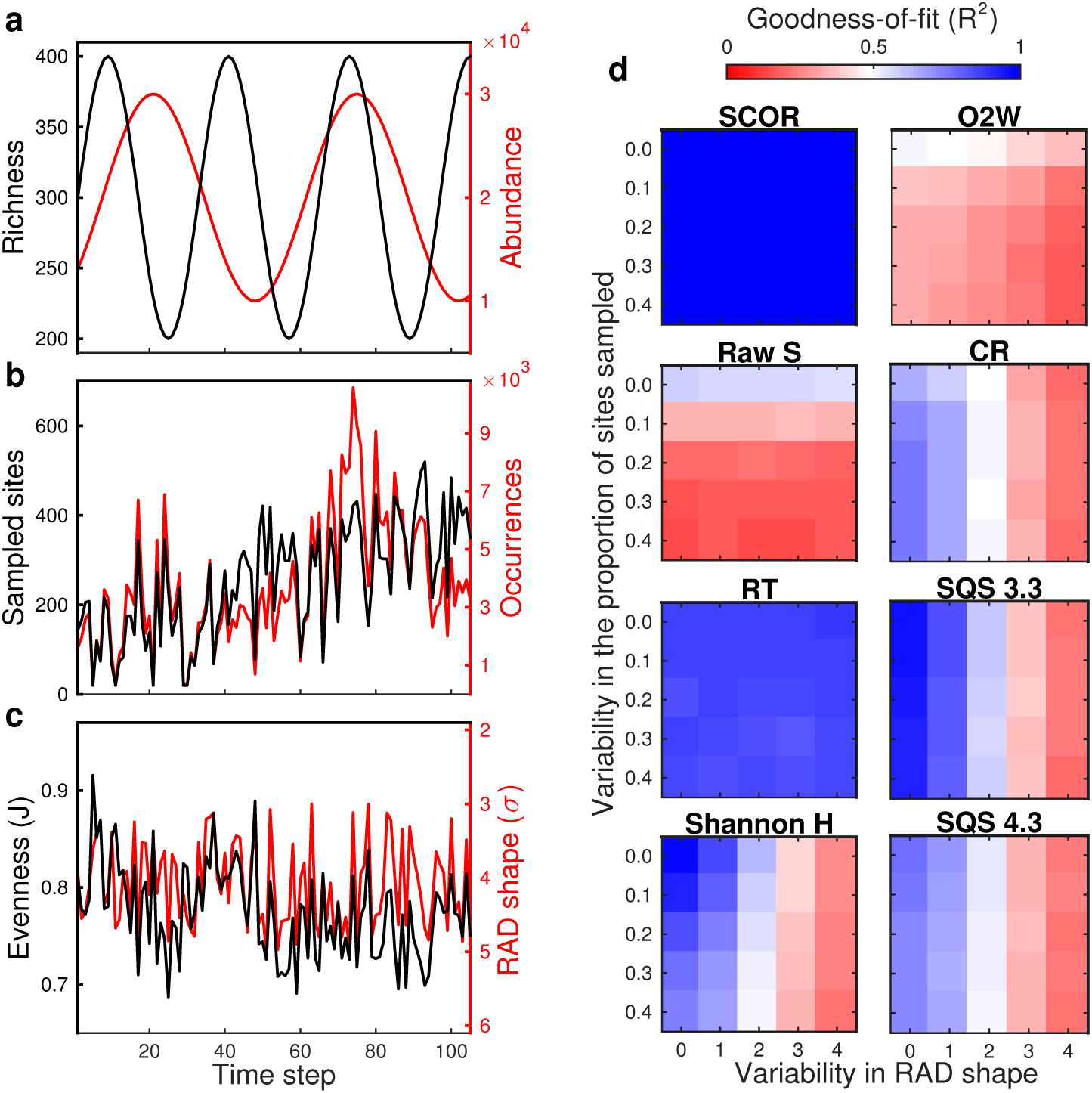
Performance of SCOR and richness estimators in Poseidon model experiments. **a**, Simulated species richness and total abundance are decoupled. **b**, Sampled species occurrences reflect abundance distorted by the trend and short-term variability in sampled sites (in this example, variability = 0.1, corresponding to the standard deviation around the mean trend). **c**, Sampled species evenness (Pielou's J) captures changes in the shape parameter σ of the RAD (in this example, variability = 2, corresponding to the range of σ), superimposed on richness fluctuations and a net decrease caused by the sampling trend. **d**, Sensitivity to sampling variability and RAD shape variability. Values are median goodness-of-fit (R^2^) of 50 model runs, comparing SCOR to true abundance, and richness estimates to true richness. See text for abbreviations.

Our simulations with Poseidon show that the SCOR estimate of relative changes in total abundance is highly robust to variability in both spatial sampling and RAD shape (Fig. 1d). By definition, SCOR is immune to the loss of rare species, and decoupled from changes in richness. As expected, the fidelity of raw sampled richness (S) decays rapidly with increasing sampling variability, but shows little sensitivity to changes in the shape of the RAD. Simple range-through richness (RT; assuming a species existed in all time bins between its first and last occurrence) is relatively robust to both factors, indicating that the level of sampling in Poseidon is sufficient to avoid severe edge effects. The Shannon entropy H,which reflects both richness and evenness, is very sensitive to RAD shape variability, ultimately tracking changes in evenness at the expense of changes in richness. Classical rarefaction (CR) and shareholder quorum subsampling (SQS)^5^, being sampling-standardization methods, are robust to the effect of spatial sampling variability on richness, all else being equal. However, both CR and SQS are highly sensitive to changes in RAD shape. As with Shannon H, increasing RAD variability causes CR and SQS to lose track of richness and respond to changes in the shape parameter σ of the RAD instead. Note that the σ values used in Poseidon generally correspond to high, moderately variable species evenness (Supplementary Fig. 1). A third subsampling method, occurrences-squared weighted (O2W)^12^, shows overall poor agreement with true richness.

Turning to the rich deep-sea sedimentary record of the Cenozoic Era (0-65 million years ago), we analysed global occurrences of the two most prominent groups of calcifying plankton, coccolithophores and foraminifera, from the Neptune Sandbox Berlin (NSB) database^13,14^ (Methods). In both groups, raw S generally increases along with the number of boreholes representing the spatial sampling, while sampled evenness (J) decreases (Supplementary Fig. 2a,b), as expected if improved sampling enhances the detection of rare species (Fig. 1b,c). Sampling-standardized richness estimates (SQS) seem to remove the sampling trend, but given the sensitivity of subsampling methods to RAD shape found in Poseidon, we suspected an evenness signal in the SQS estimates. Indeed, SQS richness can be reproduced by simply adding together the raw S and J curves (Supplementary Fig. 2c,d), a relationship that emerges across NSB data and simulation runs (Fig. 2). This result implies that changes in evenness are a major confounding factor for current sampling-standardized richness estimators.

**Figure 2.**
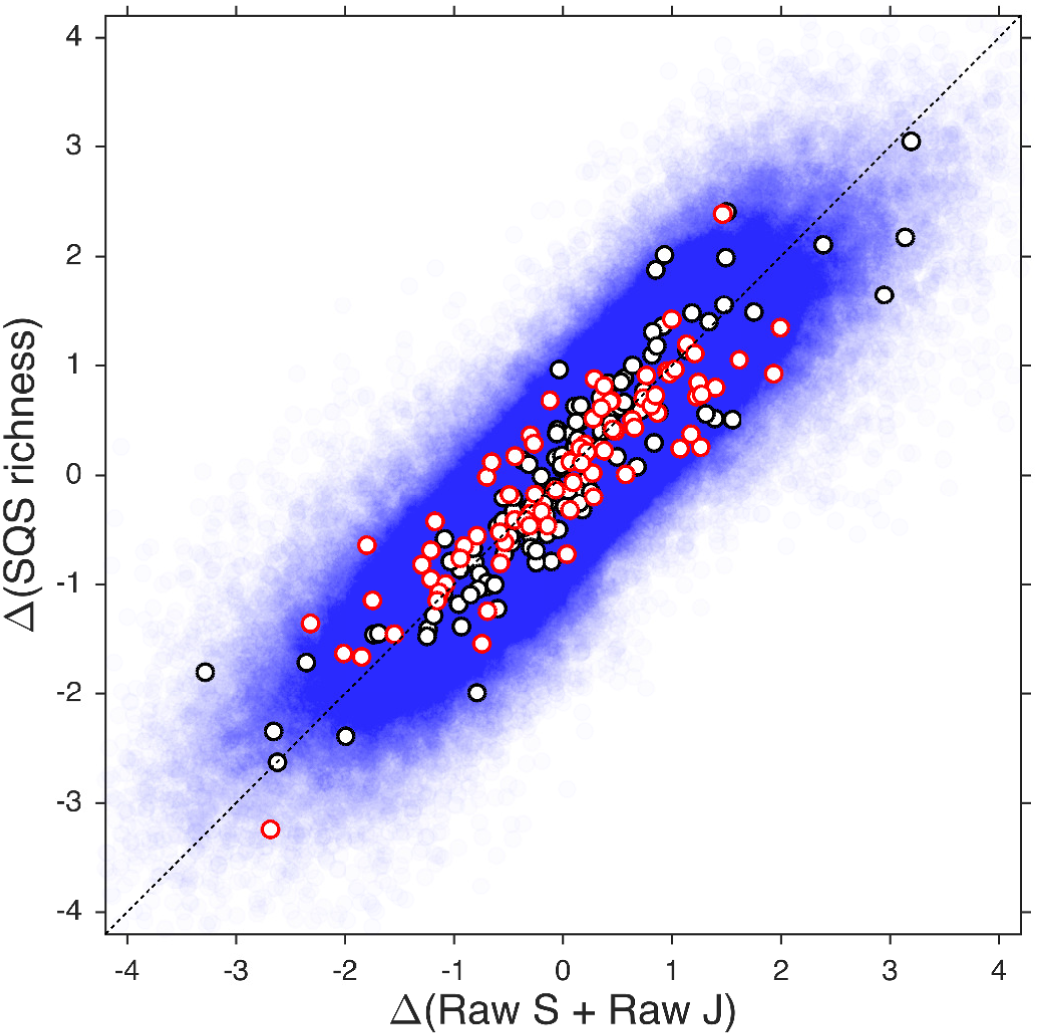
Empirical relationship between sampling-standardized richness and the sum of raw richness and evenness. Values are first differences of normalized time series of SQS richness and of the sum of normalized raw richness (S) and evenness (Pielou's J). Data include Cenozoic coccolithophores (black) and planktonic foraminifera (red) from the NSB database (Supplementary Fig. 2), and all Poseidon model experiments (blue; *N* = 262,500). Stippled line marks the 1:1 relationship.

Focussing instead on common species, coccolithophores and foraminifera have markedly different SCOR trajectories through the Cenozoic. On average, coccolithophores have their highest SCOR values in the Eocene, followed by a decline in the Oligocene and a resurgence in the late Miocene and Pliocene. Aspects of the coccolithophore SCOR pattern have been linked to Cenozoic proxy records of atmospheric CO_2_, suggesting that coccolithophores could thrive in a high-CO_2_ world^11,15^. Since their rise in the Mesozoic, coccolithophores shifted the dominant locus of carbonate burial from continental shelves to the deep sea, providing a new mechanism for buffering ocean chemistry and atmospheric CO_2_ through carbonate compensation^16^. Oligocene cooling and CO_2_ decline was accompanied by a lowering of the carbonate compensation depth, which has been attributed to changes in the supply of weathering products to the ocean^17^. The Oligocene reduction of coccolithophore SCOR is opposite to that expected if SCOR were biased upward by enhanced deep-sea preservation^11^, and carbonate preservation trends cannot explain the independent SCOR patterns in the two calcifying groups. Selective dissolution or taxonomic preferences in sample processing may cause short-term volatility in SCOR, but only if species presence or absence is random with respect to commonness (Supplementary Fig. 3).

Planktonic foraminifera SCOR was compared to Cenozoic deep-ocean temperature (DOT) records^18^ (Methods; Supplementary Data Set). Although the net trends are inversely related (foraminifera flourish as the world cools), shorter-term changes suggest positive covariation, including the Early Eocene climate optimum, Eocene cooling, as well as Miocene and Pliocene optima (Fig. 3a). Geological proxy records are generally noisy mixtures of signals representing multiple processes, derived from a sedimentary record that is itself an active component of the Earth system. Any causal connection detected between proxy records would necessarily be indirect with respect to the underlying processes of interest. Nonetheless, the DOT record reflects a set of climate-related variables, including changes in ocean thermohaline circulation, water mass structure, and nutrient dynamics, all considered to be important abiotic controls on the long-term evolution of planktonic foraminifera^6,19-21^. Here we tested this drive-response hypothesis using three conceptually very different methods for causal detection in time series (Methods): (1) Convergent cross mapping^22^, based on the concept of state space reconstruction from time-delay embedding; (2) Information transfer analysis ^23,24^, based on the concept of transfer entropy^25^; and (3) Bayesian inference of causal models based on linear stochastic differential equations^26,27^.

**Figure 3.**
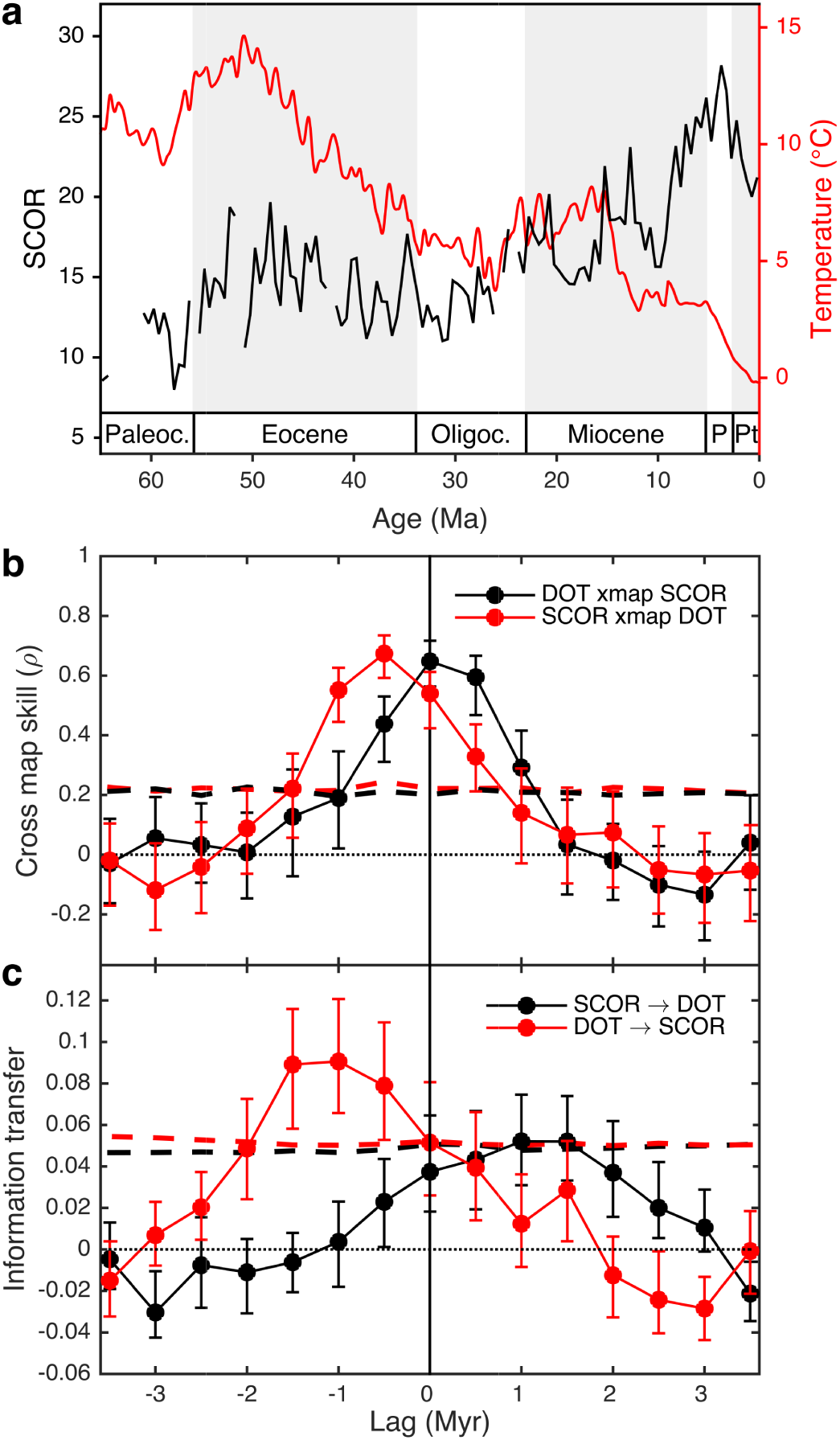
Testing a causal link between planktonic foraminifera SCOR and Cenozoic climate changes. **a**, SCOR of planktonic foraminifera from the NSB database at 0.5 Myr resolution, and DOT estimates^18^ at 0.1 Myr resolution. **b**, **c**, CCM skill (**b**) and IT (**c**) between SCOR and DOT as a function of time lag. If past DOT drives SCOR, then SCOR *xmap* DOT, while information flows DOT → SCOR, at negative lags. Values are medians (dots) and 95% ranges (whiskers) for 500 random subsamples of length 100, dashed lines are 95^th^ percentiles of 1,000 surrogates. All values are normalized to a surrogate mean of zero. Ma, million years before present; Paleoc., Paleocene; Oligoc., Oligocene; P, Pliocene; Pt, Pleistocene.

Convergent cross mapping (CCM) from foraminifera SCOR to DOT peaks at a negative lag, indicating that the SCOR signal carries a response to past changes captured in the DOT record (Fig. 3b). The optimum lag is a single time bin, implying a causal delay of 0.5 million years (Myr) or less. CCM is also significant in the opposite direction but this is stronger at positive lags (Fig. 3b), which are non-causal (future "drives" past). This result is consistent with a unidirectional forcing where the dynamics of the response variable (SCOR) is dominated by the driving variable (DOT), such that predictability flows both ways^28^. Information transfer (IT) analysis supports this inference: predictive information flow is significant from past DOT to SCOR, although the optimal lag is shifted backward by one time bin, implying a more protracted causal delay (Fig. 3c). In the opposite direction, IT peaks at the corresponding positive (non-causal) lags, but is significantly weaker than in the causal direction. Using a series of linear Stochastic Differential Equations (SDEs) to model correlation and causality between the two records (Supplementary Fig. 4), we recover relatively strong evidence that SCOR responds to changes in DOT, with a time lag of 0.3-31.1 Myr, comparable to the CCM and IT analyses (Supplementary Tables 1, 2), although the detailed nature of the causal relationship cannot be clearly resolved (Methods).

The congruence of these results strongly suggests that the ecological prominence of planktonic foraminifera has evolved in response to past climatic and oceanographic changes captured in the deep-ocean temperature proxy record. Furthermore, the inferred time delay implies that the causal connection is highly indirect, involving climate changes propagating through the Earth system to influence the commonness of foraminifera in the global plankton on evolutionary time scales. In the modern global ocean, eukaryotic plankton richness involves a vast number of parasite and symbiont species, highlighting the importance of biotic interactions in driving diversification through trophic connectivity and complexity^4^. Abiotic factors, such as differences in nutrient level among ocean basins, are more clearly reflected in the relative abundance of the dominant species. A restructuring of water masses and nutrient distributions is likely to cause a dramatic and discernible shift in the distribution and abundance of many species, yet have a far less predictable impact on richness. Our results imply that if such a fundamental ecosystem response were to leave a signature in the fossil record, it would be far more evident in the robustly detectable distribution of the most common species than in the indeterminate richness of rare species. Dominant groups also reveal macroevolutionary trends in functional morphology otherwise obscured by rare taxa^29^. Given their critical importance to ecosystem functioning, common species provide a nexus for understanding the role of an evolving biota in global environmental changes of the past.

## Acknowledgments

We thank the contributors to the Neptune Sandbox Berlin database. Thanks to J. Alroy for sharing his SQS 4.3 script. This project used DSDP/ODP samples provided by the Integrated Ocean Drilling Program (IODP). IODP is sponsored by the U.S. National Science Foundation (NSF) and participating countries under the management of Joint Oceanographic Institutions (JOI), Inc. The authors were funded by the Norwegian Research Council (231259 to B.H. and 235073 to L.H.L.) and the Bergen Research Foundation (B.H. and K.A.H.).

## Author Contributions

B.H. and L.H.L designed the study. B.H. ran the Poseidon simulations, analysed data, performed the IT analysis, and wrote the paper. L.H.L. retrieved the NSB data, calculated SCOR, and wrote the Poseidon code. K.A.H. and D.D. performed the CCM analysis. T.R. performed the SDE analysis. All authors discussed the results and commented on the manuscript.

## Additional Information

Supplementary information is available in the online version of the paper. Reprints and permissions information is available at www.nature.com/reprints.

## METHODS

### Data

Microfossil occurrences were retrieved from the NSB database^13,14^ (accessed April 22, 2015). SCOR and richness estimates were calculated using 0.5 Myr time bins. For the planktonic foraminifera, we compared the inferred times of species rise and fall in NSB^10^, which encompass any period of potential commonness, to the species ranges in the PlankRange database^30^ (http://palaeo.gly.bris.ac.uk/Data/plankrange.html, accessed Aug. 24, 2014). After resolving most of the taxonomic discrepancies, ~82 % of the species have a rise-fall interval that fits within the proposed range or is offset by < 2 Myr (a single time bin in ref. 10). Of the remaining species, which could either not be matched taxonomically or have a significant range offset, ~73 % are rare (peak occurrence frequency, as a proportion of all sites with at least one species sampled, < 0.2), and none have a peak occurrence frequency > 0.4. Taxonomic or range errors in NSB are therefore unlikely to have a significant impact on SCOR, which is sensitive only to the most widespread species. Quantification of this impact awaits the public release of the updated PlankRange database^31^.

DOT estimates were obtained from ref. 18, using their > 9 Ma T_d-SL_ record (based on subtracting New Jersey sea level records from a benthic δ^18^O stack) and scaled δ^18^O record for the interval < 9 Ma, with their interpolation at 0.1 Myr resolution. Both SCOR and DOT were tied to the GTS2004 time scale^32^.

### SCOR

We treat the observation of a specific number of individuals as a Poisson-distributed variable with parameter *λ* in each time bin. The probability of finding an individual of species *i* in time bin *j* is then *p*_*ij*_ = 1 − exp(−*λ_ij_*), and thus *λ*_*ij*_ = − ln(1 − *p*_*ij*_). In practice, *p*_*ij*_ is estimated as *y*_*ij*_/*n*_*j*_, where *y*_*ij*_ is the number of sites in which species *i* is observed at time bin *j* and *n*_*j*_ is the number of sites in that time bin where at least one of the species included in the analysis is observed. SCOR is the total density of a given set of *m*_*j*_ species in a time bin:

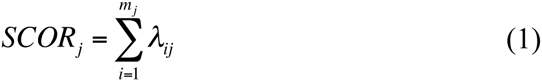

and we estimate its variance by the delta method^33^:

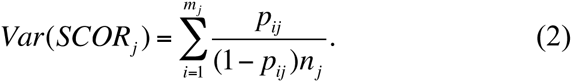

SCOR is based on the observation that the more globally abundant a species is, the more likely it is to occur at a greater number of sites^4^. As *p* approaches 1, the rate of increase in *λ* grows rapidly, such that very widespread species have a much greater influence on SCOR than restricted species. If a species occurs at all sites in a time bin, its *λ* for that time bin is undefined.

SCOR is decoupled from species richness and relative abundance. If a species becomes globally more abundant and widespread in a time interval, then its *λ*, and thus SCOR, will increase. Even if all species became exactly equally more common in absolute terms, with no change in relative abundance, their individual *λ* values will be higher and SCOR will capture the proportional change in absolute abundance of the total species set.

### Poseidon simulations

To evaluate the performance of SCOR relative to commonly used diversity metrics, we designed a set of numerical experiments on the effects of temporal variability in three factors: (1) spatial sampling completeness; (2) the shape of the species rank-abundance distribution (RAD); and (3) the proportion of species lost randomly (with respect to abundance).

Poseidon consists of 1,000 spatial grid cells and 105 time steps, where true species richness and abundance are allowed to vary independently (Fig. 1a). In each time step, we randomly assign an abundance value to each species, such that the entire community has a log-normal rank-abundance distribution (RAD), the shape of which can be fixed or time-varying. Species ranks are randomly reshuffled between time steps. We then randomly assign a spatial grid cell (site) to an individual of a species, where it can potentially be preserved and sampled.

Next, we sample only a proportion of the sites (spatial cells) such that this proportion increases linearly from 0.1 to 0.4, representing a declining sampling coverage with age, typical of deep-sea sedimentary records. Any short-term variability is thus superimposed on this trend (e.g. Fig. 1b). Furthermore, a proportion of the remaining species can be randomly removed (representing dissolution, selective picking, or other processes causing a species to be absent in a time bin, regardless of its original abundance).

We then calculate raw S, RT, Shannon H, and three sampling-standardized richness metrics widely used in palaeobiology (CR, O2W, and SQS). Although a generalized OXW has been recommended for paleontological datasets^34^, we use O2W here because our data meet the assumptions of the latter^12^. We used two versions of the shareholder quorum subsampling (SQS) method^5,35-37^: the SQS3.3 R script (http://bio.mq.edu.au/~jalroy/SQS.html, downloaded Aug. 26, 2014), and the SQS4.3 perl script, kindly provided by J. Alroy (Sept. 3, 2014). We did not modify the SQS codes but wrote a function to format Poseidon output to species occurrence data for SQS4.3. For SQS3.3, we input the list of sampled individuals (abundances). A quorum level of 0.6 was used in all runs. Both SQS versions yielded very similar results when given the same type of data (abundances or occurrences). For all subsampling methods (CR, O2W, and SQS), 100 iterations were used in obtaining each estimated richness value. Increasing the number of iterations offered no discernible improvement. Shannon H was output from the SQS 3.3 R script and used to calculate Pielou's J evenness. The goodness-of-fit between true and estimated richness, and between true abundance and SCOR, was assessed by the coefficient of determination (R^2^) between time series. Poseidon R scripts are provided as Supplementary Code.

### Time Series Analysis

We used three different time series analysis methods to test for a causal relationship between SCOR and DOT. Two of the methods are non-parametric, while the third is based on linear models. To implement a time-displacement (lag) analysis for the non-parametric methods, missing SCOR values were linearly interpolated, and the 0.1-Myr resolution DOT record was bin-averaged on the SCOR time bins (0.5 Myr). Furthermore, the non-parametric methods use surrogate time series to assess significance, which requires detrending of the original time series. To avoid any bias from differences in non-stationarity that are not reproduced by the surrogates, both records were detrended using a third-order polynomial to satisfy a stationarity criterion^38^, then normalized to zero mean and unit standard deviation. For consistency, the model-based analysis was also performed on the preprocessed data. However, the detrending may remove components of the variation relevant to uncovering the parameters of underlying processes. We therefore repeated the model-based analysis on untransformed data.

### CCM analysis

CCM is a method for causal inference in nonlinear dynamical systems based on the theory of state-space reconstruction^22^. Consider two time series *X* and *Y* consisting of scalar observations of variables in a dynamical system. According to Takens's theorem^39^, we can construct a delay-coordinate embedding of the state space of the dynamical system into an m-dimensional real space, by constructing the vectors *E*_*X*_ = { ⟨(*x*(*t*_i_), *x*(*t*_i_ – τ), *x*(*t*_i_ – 2τ), …, *x*(*t*_i_ – (*mτ*) ⟩}, where *x*(*t*_i_) is the scalar value of the time series *X*at time *t*_i_. That is, the vectors in *E*_*X*_ are in one-to-one correspondence with the states of the dynamical system. If *X* and *Y* are coupled variables of the same dynamical system (i.e. they are causally influencing each other), this correspondence is also true for time series *Y*, and therefore *E*_*X*_ and *E*_*Y*_ are in one-to-one correspondence with each other. CCM uses this result to predict scalar values of *Y* from the coordinate-delay embedding of *X* and vice versa.

The CCM algorithm locates, for each scalar point *P*_i_ in the prediction set (subset of time series *Y*), the contemporaneous state vector *L*_i_ in the library set (subset of state vectors in the time-delay embedding constructed from time series *X*). Next, it finds *L*_i_'s closest neighbours and estimates a value for the predictee *P*_i_* using simplex projection^40^. CCM skill is determined by the correlation (Pearson's ρ) between estimated *P*_i_* and actual values of *P*_i_.

With increasing library size, CCM skill is expected to increase and converge if the variables are causally related. The notation “*X* xmap *Y*” refers to estimating *y*(*t*_i_) from corresponding lagged-coordinate state versions of *x*(*t*_i_), which in a causal context is read as “*Y* is causally influencing *X*”.

CCM analysis was performed using the rEDM software package^41^. We constructed time-delay embeddings using embedding dimension *m* = 2 and delay time step *τ* = 1. Cross mapping was then performed using the entire time series as both prediction and library sets. To avoid biased results, we used leave-one-out cross validation (i.e. the predictee *P*_i_ itself and points in a time radius of *E* around *P*_i_ were excluded from the libraries, such that no points sharing coordinates with *P*_i_ were used in the predictions; see refs. 41, 42).

If unidirectional forcing is sufficiently strong, the dynamics of the response variable can become dominated by the driving variable. In this case, CCM may be significant in both directions, and thus unable to distinguish unidirectional forcing from bidirectional causality. To address this, we used the extended CCM approach ^28^, which repeats the cross mapping using different time-displacements of the original time series. If there is a discernable lag between cause and effect, then optimal cross map skill is expected to occur for negative time lags in the direction(s) of true causality (past drives future). If true causality is unidirectional, then any CCM skill in the non-causal direction is expected to peak for positive lags (future "drives" past).

Extended CCM analysis of SCOR and DOT is reported as median cross map skill and 95 % ranges at different lags after drawing 500 samples with replacement from libraries of size 100. Statistical significance is evaluated against a null distribution of CCM results for 1000 surrogate time series. For each lag, CCM skill is considered significant if it exceeds the 95^th^ percentile of the surrogates. We verified the results using three different methods for generating surrogate data: phase-randomized and amplitude-adjusted Fourier transform (AAFT)^43^, phase-randomized Fourier transform^44^, and randomly shuffled surrogates. All three methods indicate significant causality from DOT to SCOR, and we limit our results to the AAFT method, which gave the most conservative significance estimates.

### IT analysis

If two processes *X* and *Y* are independent, then a general Markov property will hold^25^:

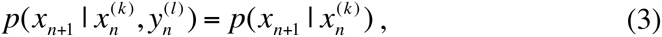

where *p(*x**_*n*~1_) is the transition probability to state *n+*1, and indices *k* and *l* are the dimensions of vectors of past states. In the absence of information flow from *Y* to *X*, knowing the past *l* states of *Y* has no influence on the transition probability of *X* beyond knowing the past *k* states of *X* alone. Transfer entropy^25^ quantifies the incorrectness of assuming independence by means of a Kullback-Leibler divergence, a non-symmetric measure of the information lost when the right hand side is used to approximate the left hand side of equation (3):

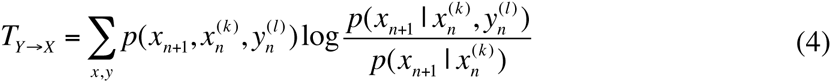

Transfer entropy is thus a non-symmetric measure of information flow, which has been shown to be equivalent to a conditional mutual information^45^, and equivalent to Granger causality^46^ for linear, Gaussian systems^47^. We implement it in the modified IT form proposed by Verdes^23^, which has previously been applied to the analysis of geological records^11,24,48,49^.

Here we expand on these earlier applications by repeating the IT analysis for different time-displacements of the original time series, analogous to the extended CCM analysis described above. Similar to CCM, predictive information flow may become symmetric if unidirectional forcing and/or linear correlation is sufficiently strong. However, if there is a discernable lag between cause and effect, then optimal information transfer is expected to occur for negative time lags in the direction(s) of true causality (past → future). If true causality is unidirectional, then any information flow in the non-causal direction is expected to peak for positive lags (future → past). IT is a coarse-grained relative entropy measure, which varies as a function of the data gridding resolution, summarized in a single IT value as the area under the resulting curve^23^. Lagged IT analysis of SCOR and DOT is reported as median IT and 95 % ranges at different lags after drawing 500 random subsamples of size 100. IT is considered significant if it exceeds the 95^th^ percentile of a null distribution of IT results for 1,000 AAFT surrogate time series. Unlike CCM, this IT implementation does not use time-delay embedding. Combined with coarse-graining of the dat**a**, this may help explain the difference in optimal lag between IT and CCM (Fig. 3b,c), although more work is needed to clarify this.

### Linear SDE analysis

Given two time series representing two measured processes, linear SDEs can be used to distinguish between correlation and Granger causality. Uni-and bidirectional causation as well as hidden (unmeasured) processes can be modelled in the SDE framework, expanding the space of possible connections ^26,27^ A basic linear SDE can be written as

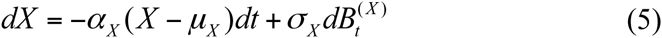

This describes a mean-reverting Ornstein-Uhlenbeck process (OUP) *X*, which contains a systematic part (the *dt* term) and a stochastic part (*t*he *dB* term). If the systematic part is dropped (*α*_*X*_ = 0), then equation (5) describes a Wiener process (WP, or random walk). The OUP has expectation *μ*_*X*_, stationary standard deviation 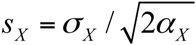 and half-life *t*_1/2,*X*_ = log(2)/ *α*_*X*_. To model a hidden process, we can write

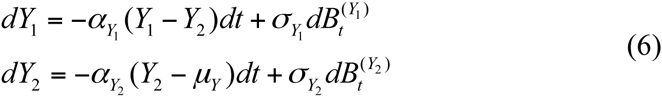

Here, the measured process *Y*_*1*_ has a hidden process (or layer) *Y*_*2*_ folded into its systematic part, such that *Y*_*1*_ tracks *Y*_2_. *Y*_*1*_ is similar to an OUP, but instead of fluctuating around a fixed expected value it fluctuates with a lagged response to the OUP *Y*_2_.

When modelling connections between processes, we use vector notation. A pure correlation between *X* and *Y* entails that the covariance matrix in front of the stochastic term *d<u>B</u>* will have off-diagonal elements. If there is a causal connection from *Y*_*2*_ to *X*, for instance, the system takes the following form

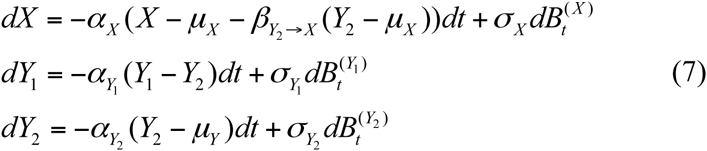

where *β*_*Y*_2__ → *X* describes the connection strength from *Y*_*2*_ to *X.* Equation (7) describes a “common cause” situation, where *Y*_*2*_ drives both *X* and *Y*_*1*_.

To analyse the SCOR and DOT records, we first characterized each time series separately, examining models with up to three layers (two hidden). In each model, the layers could be WP or OUP (including fully deterministic layers where *σ*_*i*_ = 0), excluding a one-layered WP, which prohibits incoming causal links. We also excluded internal feedback loops in multi-layer models because of numerical intractability. For both time series, *μ* was assigned a prior distribution *μ*_*i*_ ~ *N*(0,1), where *i* denotes the layer. All model parameters were assigned normal priors, with 95 % prior probability ranges of *σ*_*i*_ ∈ (0.01,1.0) for the stochastic term, *t*_1/2_ ∈ (0.1*My*,50*My*) for the half life, *β*∈ (-2,2) for the causal connections, and *ρ*∈ (-0.96,0.96), for the logit-transformed correlation coefficients. We used MCMC importance sampling to estimate Bayesian model likelihoods and calculate model probabilities.

The best model for SCOR in isolation was a one-layered OUP, while a three-layered model with a WP as the bottom driver was the best model for DOT in isolation. We then investigated all 15 connection models between these two best models, including the null hypothesis of no relationship (Supplementary Fig. 4). We allowed for causality from SCOR to DOT because both proxy records ultimately derive from deep-sea carbonate sediments, hence SCOR could in principle contain a signal of processes that have influenced DOT. The null hypothesis was assigned 50 % prior probability, while 50 % was distributed evenly among the 14 connection models. For model comparison, we used Jeffreys’s scale to assess the strength of evidence represented by the Bayes factor B, where 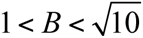 is evidence “barely worth mentioning”, 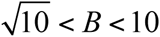 is “substantial evidence”, 10 < *B* < 10^3/2^ is “strong” evidence and *B* < 10^3/2^ is “very strong” evidence^50^.

The posterior probability of the null hypothesis was 11.2 % (Model 1; Supplementary Fig. 4), hence the Bayes factor favouring a connection between SCOR and DOT is 7.9 (substantial evidence). The most probable model (Model 5; Supplementary Fig. 4) involves a feedback loop, where the upper DOT process (DOT1) affects SCOR positively while SCOR affects DOT1 negatively. All parameter estimates with credible intervals for the best model are presented in Supplementary Table 1. In the second most (and almost equally) probable model, (Model 12; Supplementary Fig. 4), SCOR affects the second DOT layer (DOT2) instead.

With a half-life of ~0.5 Myr, SCOR responds to DOT processes on time scales comparable to those inferred from the other analyses (Fig. 3b,c). In contrast, DOT processes react very slowly to changes in SCOR (Supplementary Table 1), and because SCOR changes rapidly, the response in DOT will be smoothed out. From equation 5, the effect of SCOR on DOT1 is 0.07 Myr^-1^, while the effect of DOT1 on the SCOR process is 0.6 Myr^-1^. Thus, DOT influences SCOR much more strongly per time unit than vice versa.

The third most likely model (Model 4; Supplementary Fig. 4) only had a causal connection from DOT1 to SCOR, consistent with CCM and IT inferences. This model had a posterior probability of 17.5 %, hence the evidence for a feedback loop is less than “substantial" (B = 2.6). In summary, we find evidence for there being at least one connection (B = 7.9); for the connections to be causal rather than correlative given that there are connections (B = 12.5); and specifically for a causal connection from DOT to SCOR given that there are connections (B = 15.4).

We then repeated the analysis on untransformed data (not detrended or normalized), denoted uDOT and uSCOR. In this case, the best isolated model for both time series is a three-layer model with a WP at the bottom. The Bayes factor favouring a connection over no connection is 73, which is deemed “very strong evidence”. The best connection model involves a feedback loop between the top layers uDOT1 and uSCOR1. However, there is a very high probability for parameters enforcing cyclical behaviour, with a period of 1.5 Myr, which is consistent with an internal feedback loop model as the best isolated model for uDOT. Parameter estimates are shown in Supplementary Table 2.

---

**Supplementary Figure 1.**
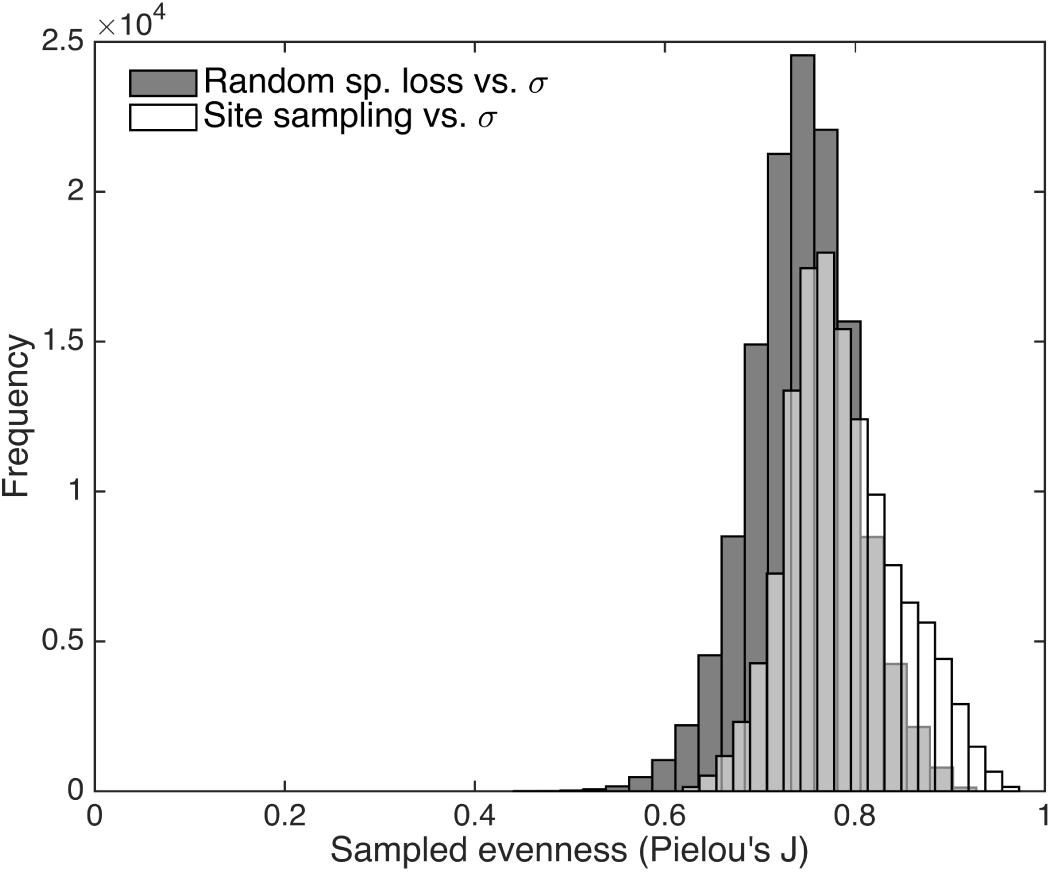
Distribution of sampled evenness across all Poseidon experiments. Shaded histogram represents the model runs testing the sensitivity to variability in the proportion of species randomly removed, and variability in RAD shape parameter σ (Supplementary Fig. 3). Un-shaded histogram (note transparency in overlap) represents the model runs testing the sensitivity to variability in the proportion of sites sampled, and variability in σ (Fig. 1). The median evenness is 0.76, and 95 *%* of the values are in the range 0.65 – 0.90.

**Supplementary Figure 2.**
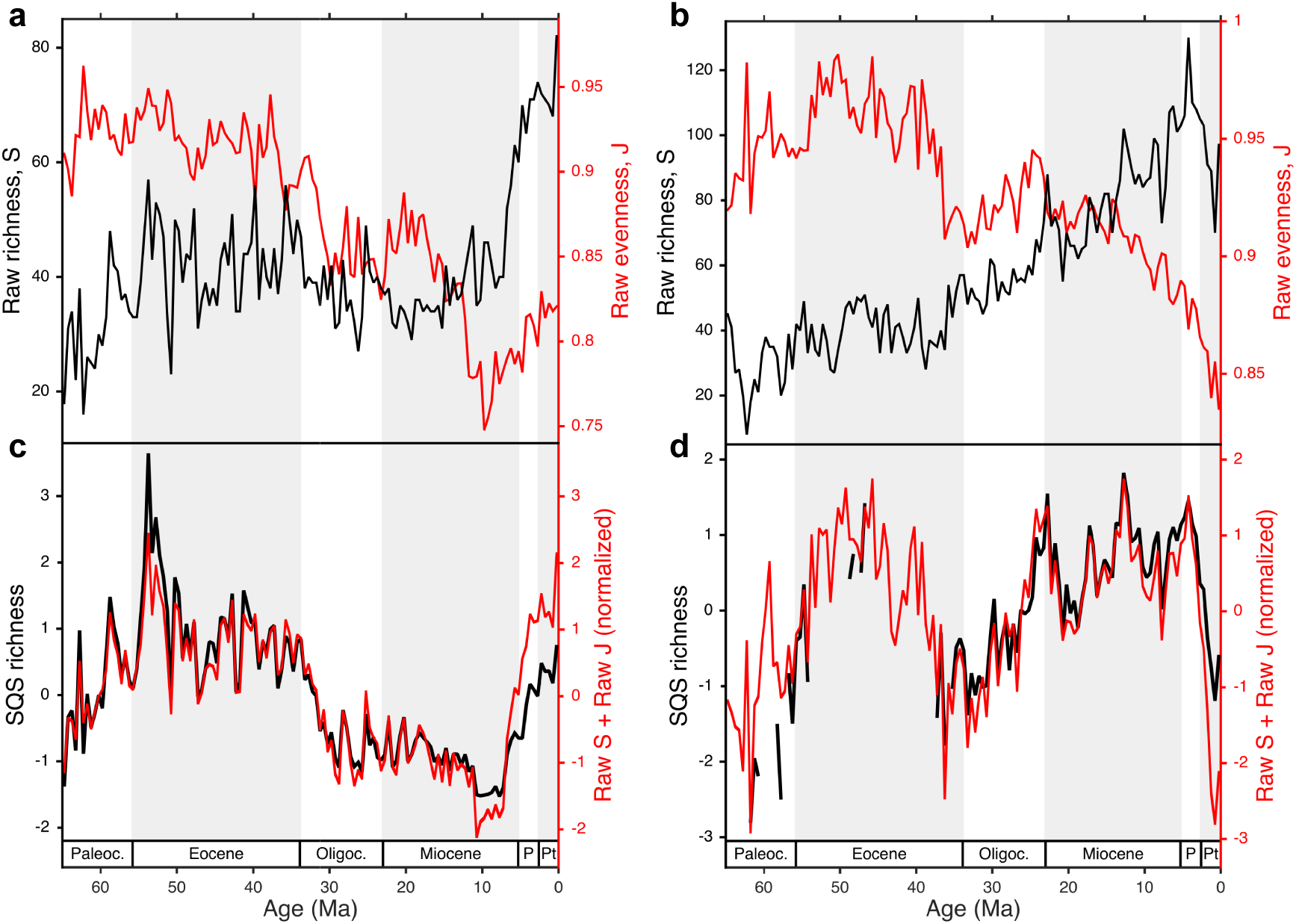
Sampling-standardized richness can be reproduced by the sum of raw richness and evenness. **a**,**b**, Raw sampled richness (S) and evenness (Pielou's J) of Cenozoic coccolithophores (**a**) and planktonic foraminifera (**b**) species from the NSB database. **c**, **d**, The sum of raw S and raw J superimposed on shareholder quorum subsampling (SQS) estimates of richness for coccolithophores (**c**) and foraminifera (**d**), all normalized to zero mean and unit standard deviation. SQS was calculated with a quorum level of 0.4 (higher quorum levels give nearly identical results but are less complete for the older part of the record). Ma, million years before present; Paleoc., Paleocene; Oligoc., Oligocene; P, Pliocene; Pt, Pleistocene.

**Supplementary Figure 3.**
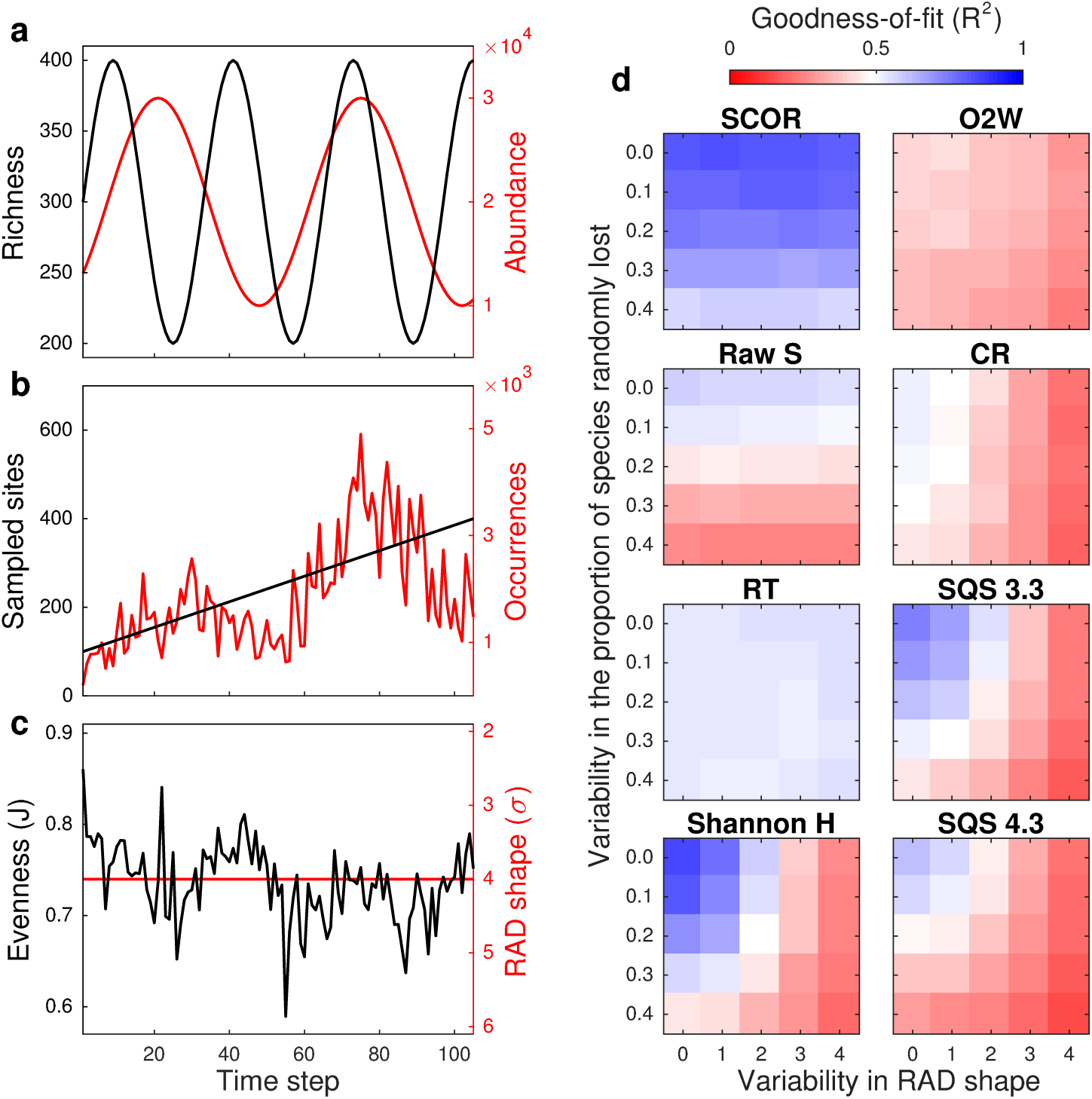
Effect of random species loss in Poseidon model experiments. **a**, Simulated richness and abundance as in Fig. 1a. **b**, Site sampling increases smoothly in all experiments. Instead, a proportion of the species is randomly removed in each time step, causing volatility in occurrences. No variability in the proportion lost means that 50 *%* are always removed. In this example, variability = 0.4, meaning that between 30 % and 70 % of species are lost. **c**, Even with a constant original RAD shape, random species loss, and variability in the proportion lost, generates volatility in sampled evenness (*t*his example is an extreme case, see Supplementary Fig. 1). **d**, Sensitivity to variability in RAD shape and in the proportion of species lost. Values are median goodness-of-fit (R^2^) of 50 model runs, comparing SCOR to true abundance, and richness estimates to true richness.

**Supplementary Figure 4.**
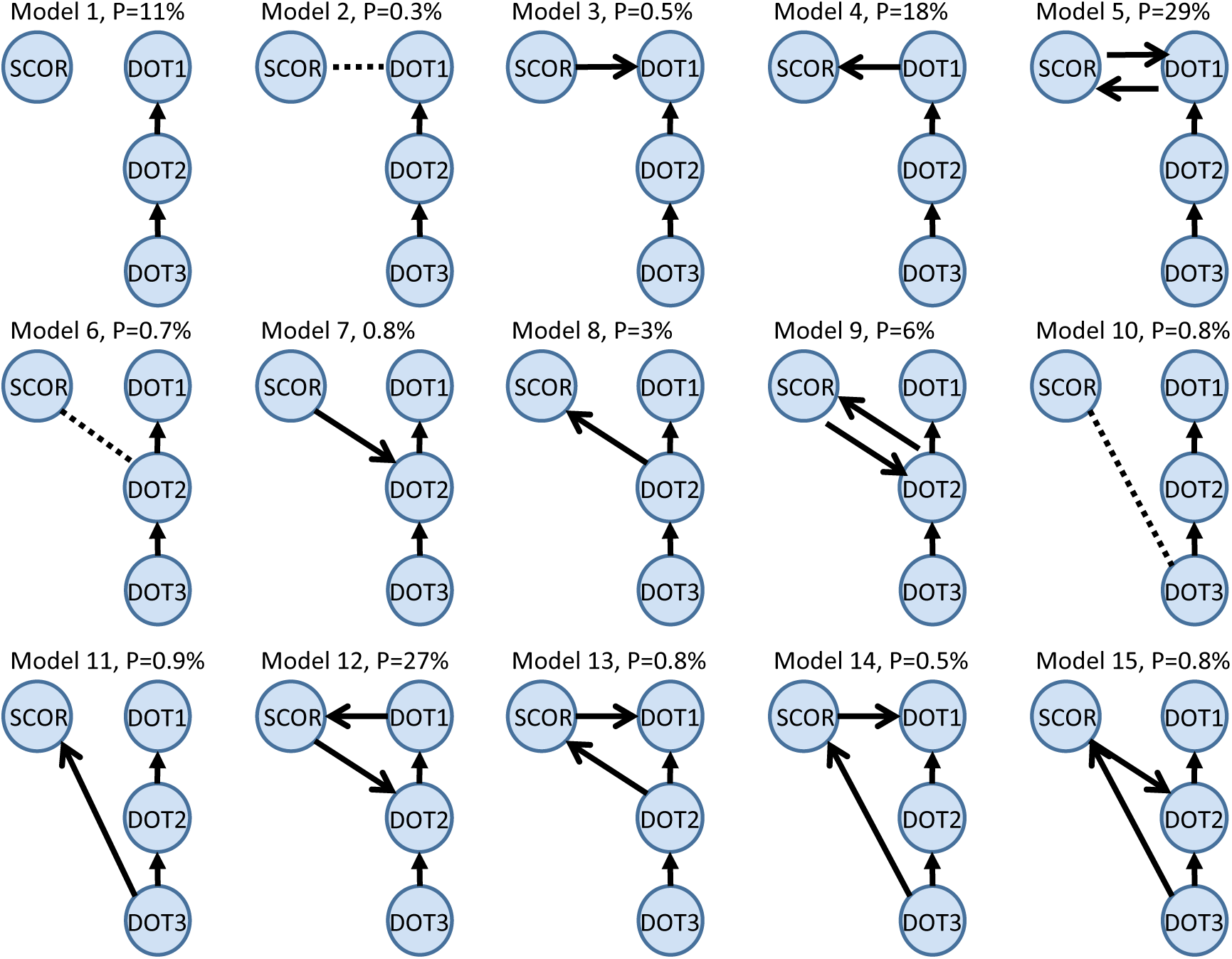
Schematic of all connection models between SCOR and DOT in linear SDE analysis. The best model for SCOR in isolation is a one-layered OUR The best model for DOT in isolation is a three-layered model with a WP as the bottom layer (DOT3). All models possible between these two best models are shown. Note that SCOR cannot drive DOT3 because DOT3 is a WP. Percentage values represent posterior model probabilities. Solid arrows represent casual connections pointing from driver to response. Dotted lines represent correlative relationships. See Methods for details.

**Supplementary Table 1.** Parameter estimates for the most probable connection model between SCOR and DOT.

| Parameter | Estimate (posterior median) | 95 % credible interval |
| --- | --- | --- |
| $t_{1/2,SCOR1}$ | 0.53 Myr | (0.33, 1.1) Myr |
| $s_{SCOR1}$ | 0.94 | (0.65, 1.1) |
| $\mu_{SCOR}$ | 0.43 | (-0.01, 0.94) |
| $t_{1/2,DOT1}$ | 8.0 Myr | (0.91, 31) Myr |
| $s_{DOT1}$ | 1.2 | (0.37, 2.4) |
| $t_{1/2,SDOT2}$ | 16 Myr | (3.4, 166) Myr |
| $s_{DOT2}$ | 0.45 | (0.03, 2.8) |
| $\sigma_{DOT3}$ | 0.027 | (0.003, 0.22) |
| $\beta_{DOT1 \rightarrow SCOR1}$ | 0.45 | (0.14, 0.76) |
| $\beta_{SCOR1 \rightarrow DOT1}$ | -0.77 | (-2.3, 0.5) |

**Supplementary Table 2.** Parameter estimates for the most probable connection model between uSCOR and uDOT.

| Parameter | Estimate (posterior median) | 95 % credible interval |
| --- | --- | --- |
| $t_{1/2,uSCOR1}$ | 0.91 Myr | (0.32, 1.6) Myr |
| $S_{uSCOR1}$ | 0.07 | (0.01,0.35) |
| $t_{1/2,uSCOR2}$ | 2.4 Myr | (0.86, 68) Myr |
| $S_{uSCOR2}$ | 0.7 | (0.02,21) |
| $\sigma_{uSCOR3}$ | 2.8 | (0.04, 5.8) |
| $t_{1/2,uDOT1}$ | 0.24 Myr | (0.13, 0.33) Myr |
| $t_{1/2,uDOT2}$ | 0.50 Myr | (0.21, 1.5) Myr |
| $\sigma_{uDOT3}$ | 4.1 | (1.9, 8.9) |
| $\beta_{uDOT1 \rightarrow uSCOR1}$ | 2.7 | (-2.6, 3.8) |
| $\beta_{uSCOR1 \rightarrow uDOT1}$ | -2.7 | (-5.2, 1.9) |

